# Low impact of internal stem decay on forest carbon stocks in fire-prone *Pinus ponderosa* forests

**DOI:** 10.64898/2026.05.17.725735

**Authors:** Markus Hauck, Ganbaatar Batsaikhan, Germar Csapek, Steffen Rust, Harold S.J. Zald, Choimaa Dulamsuren

## Abstract

Large old trees are of eminent importance for organic carbon storage in forest ecosystems and thus play a role in mitigating climate change. Such trees also have an increased risk of internal stem decay and tree cavity formation, which promotes biodiversity, but complicates the prediction of their biomass and carbon stocks, which is usually done from stem diameter and tree height data applying allometric biomass functions. Since the extent of internal stem decay is known to vary widely between different forest ecosystems and data from moist temperate forests exhibited low significance of internal stem decay, we studied dry, frequently fire-exposed *Pinus ponderosa* forests in central Oregon to capture the other climatic extreme of temperate forests. We hypothesized high significance of internal stem decay for stand aboveground tree biomass, as we assumed widespread stem injury from fire. In addition, we tested the hypothesis that far more than the largest 1% of trees are necessary for 50% stand biomass, as this hypothesis is found in the literature, but has been challenged in other studies. We found low biomass loss due to internal stem decay by only ca. 1% suggesting that also for fire-prone temperate forests of western North America, biomass estimates based on allometric regression are reliable. The ‘1% largest trees-50% stand aboveground biomass’ hypothesis has to be rejection for our forests as long as only trees of a size are included that noteworthily contribute to stand biomass. This metrics strongly depends on regeneration density, which is not relevant for stand biomass.

## 1. Introduction

Large old trees are long since valued for their high significance for forest biodiversity (Lindenmayer & Lawrence 2017) and their global decline has been pointed out (Lindenmayer et al. 2012). One important cause of the high value for biodiversity is the structural richness of these trees that includes tree cavities (Remm & Lõhmus 2011; Sillett et al. 2025). More recently, the great importance of large old trees for forest carbon stocks has been emphasized, as the amount of carbon stored in the biomass simply depends on tree size (Slik et al. 2013; Thom & Keeton 2019; Bordin et al. 2021; Brazee & Burcham 2023). Lutz et al. (2012, 2018) even quantified that the 1% largest trees of a forest would contain approximately 50% of the forest’s organic carbon in the aboveground tree biomass. If so much of the organic carbon stock is stored in a very small proportion of the total tree individuals of a forest stand, this provides additional support that large old trees should be kept in managed forest stands.

However, this high significance of large old trees for forest carbon estimates also raises the question if these estimates are sufficiently accurate, because biomass losses in the interior of the trees due to internal stem decay and the formation of tree cavities are routinely not incorporated in tree biomass estimates (Nogueira et al. 2006). Rather, tree biomass is usually estimated based on stand survey data of stem diameter at breast height (dbh) and tree height measurements that are used as input variables in allometric regression functions (Chojnacky et al. 2014). If the 1% largest trees represent ca. 50 % of the stand aboveground biomass, estimation errors for these tree individuals due to neglecting internal stem decay can result in significant errors for estimates of stand biomass. Even attempts were made to model stand biomass only from the biomass of the largest tree individuals (Xiong et al. 2023). The potential bias by internal stem decay was also corroborated in a modeling approach with more than 1200 trees from eastern North America, where Frank et al. (2018) found that thick-stemmed trees (expressed as the ratio of dbh to tree height) and tall trees were more strongly subjected to internal stem decay than smaller trees.

The topic that internal stem decay can reduce the biomass of individual trees significantly has been addressed in the literature repeatedly (Zheng et al. 2016; Marra et al. 2018; Wei et al. 2022; Aishan et al. 2024), but rarely been quantified with respect to effects on the stand level, which would have the potential to influence the estimates for entire biomes, countries or continents during upscaling. Tropical savannas with high populations of termites have been identified to be subjected to substantial biomass losses due to internal stem decay on both the tree and the stand level (Calvert et al. 2024; Yatsko et al. 2025a, b). On the stand level, aboveground biomass losses due to internal stem decay of up to 36% were found in a dry savanna of northeastern Australia (Flores-Moreno et al. 2024). In moist tropical and subtropical rainforests, published data vary in a wide from 0.7% (Nogueira et al. 2006) via 1.7% (Clark & Clark 2000) and 3% (Flores-Moreno et al. 2024) to 8% (Zheng et al. 2016) of aboveground biomass losses due to internal stem decay and tree cavities. In boreal forests of Mongolia, internal stem decay resulted in overestimation of aboveground biomass by 4% (Dulamsuren et al. 2026). In managed temperate forests of Central Europe aboveground biomass overestimation due to internal stem decay was considerably less important, amounting only to 0.2–0.6% in various forest types dominated by *Abies alba, Fagus sylvatica, Picea abies, Pinus sylvestris*, or *Quercus robur* (Hauck et al. 2023, 2025).

The wide range of aboveground biomass losses by internal stem decay from <1% in managed temperate forests of Europe and >35% in termite-infested savannas of Australia leaves plenty of room to speculate about the global significance of internal stem decay as a potential source of error in allometry-based forest biomass and carbon stock estimates. The only way forward to fill this gap at the present state of knowledge is through case studies. Based on the studies from managed temperate forests under moist Central European climate (Hauck et al. 2023, 2025), we dedicated the present study to the effect of internal stem decay on aboveground stand biomass and carbon stocks at the dry end of the global temperate forest biome in a fire-prone forest ecosystem of western North America (Cova et al. 2025). We therefore selected forests of *Pinus ponderosa* of western Oregon, which contrast with the situation of the forests studied in Central Europe (Hauck et al. 2023, 2025) by being exposed to high fire risk during the dry summers of the western North American temperate climate (Trouet et al. 2009) and experiencing fire return intervals of few decades depending on human intervention and topography (Weisberg & Swanson 2003; Merschel et al. 2018). Forest fires are well-known to increase the risk for internal stem decay and the formation of tree cavities (Adkins 2006; Bär et al. 2019). Like most forests of the temperate zone, also most *P. ponderosa* forests are managed for both timber harvest and fuel reduction (Hagmann et al. 2013; Westlind & Kerns 2017; Merschel et al. 2021; Fettig et al. 2025). Detecting tree cavities with sonic tomography, we tested the hypothesis that biomass losses in this dry, fire-prone temperate forest ecosystem due to internal stem decay are a significant source of error in biomass estimates obtained by allometric regression from dbh and tree height data.

In addition to the quantification of biomass loss due to internal stem decay, our second objective was to examine the importance of large old trees for stand aboveground biomass density itself, testing the hypothesis by Lutz et al. (2018) that the 1% largest trees would represent ca. 50% of the stand aboveground biomass. Hauck et al. (2023, 2025) could not confirm this hypothesis in managed temperate forests under the moist climate of Central Europe, where the largest 10–25% trees were needed for 50% of the aboveground stand biomass. In an existing biomass study from *P. ponderosa* forests of Oregon, Mildrexler et al. (2020) found the 3% largest trees to constitute 42% of the stand biomass. We, therefore, hypothesized that, despite expecting a disproportionally high share of the aboveground biomass in large old trees, the ‘1% largest trees-50% stand aboveground biomass’ assumption would not be valid in the *P. ponderosa* forests. Since Hauck et al. (2023) pointed out that the percentage of trees, which is needed to sum up to 50% biomass is highly dependent on the lower size from which on young trees are included in the sample, we consistently worked with two subsets of trees, one with dbh ≥1 cm (as in Lutz et al. 2018) and another one with dbh ≥10 cm to capture trees that exert a significant effect on stand biomass.

## 2. Materials and Methods

### 2.1. Study area

Managed forests of *Pinus ponderosa* P. Lawson & C. Lawson were studied in Pringle Falls Experimental Forest located in the La Pine basin of western central Oregon (USA). Pringle Falls Experimental Forest and the neighboring Deschutes National Forest are situated on the eastern flank of the Cascade Mountains in predominantly plain, but highly elevated terrain at elevations ca. 1200–1400 m a.s.l. Geologically, the La Pine basin is dominated by deposits of sand and silt sediments and volcanoes that have been partly active during the Holocene in nearby mountain sites. La Pine (43°41’N, 121°30’W, 1290 m a.s.l.) has temperate climate with a mean annual temperature of 7.5 °C. Precipitation (ca. 500 mm a^−1^) is relatively low due to the position of the La Pine basin in the rain shadow of the Cascade Mountains and is characterized by a dry phase during summer that causes a fire season from peak summer to early autumn. Mean January precipitation amounts to 75 mm, in contrast to 11 mm in July. The *P. ponderosa* forests were strongly dominated by this species and only occasionally intermixed with small quantities of *Pinus contorta* Bol. and *Abies grandis* (Douglas ex D. Don) Lindl.

### 2.2. Sampling design

We sampled 25 plots of 400 m2 in *P. ponderosa* forests (43.709–43.805 °N, 121.586–121.660 °W) and deliberately selected stands with large old trees (Table 1). Our sample plots had a mean elevation (±standard deviation, SD) of 1333±40 m a.s.l. and covered a range of elevations from 1300 to 1460 m a.s.l. *Pinus ponderosa* stands for our study were selected based on stand survey data owned by Pringle Falls Experimental Forest (U.S. Department of Agriculture Forest Service). Because internal stem decay and tree cavity formation are most likely to occur and to influence stand biomass stocks in stands with particularly large old trees, we selected stands with a share of such trees and did not select trees of random age for our study, as we did in similar studies from other areas (Hauck et al. 2023, 2025). The design for the search of sample plots implies that our estimates of stand biomass and organic carbon stock densities are likely to exceed the average of western North American *P. ponderosa* stands. Otherwise, we had no sampling bias for certain site conditions. We also did not discriminate for certain regeneration densities during plot search, which widely varied between the *P. ponderosa* stands of Pringle Fall Experimental Forests, mostly due to differences in the fire regime caused by natural wildfire or prescribed burning.

**Table 1.** Characteristics of *Pinus ponderosa* stands.

| Parameter<sup>a</sup> |  |
| --- | --- |
| Stand density (trees ha<sup>-1</sup>): |  |
| All trees, dbh ≥1 cm | 676±173 (100 – 3275) |
| Trees, dbh ≥10 cm | 219±30 (100 – 625) |
| Trees, dbh ≥1 cm, <10 cm | 457±148 (0 – 2650) |
| Stand basal area (m ha<sup>-1</sup>): |  |
| All trees, dbh ≥1 cm | 58.30±2.56 (38.28 – 87.99) |
| Trees, dbh ≥10 cm | 57.32±2.56 (38.14 – 87.99) |
| Trees, dbh ≥1 cm, <10 cm | 1.08±0.38 (0.0 – 7.75) |
| Mean dbh (cm)<sup>b</sup> | 60.5±4.3 (26.2 – 93.0) |
| Maximum dbh (cm)<sup>b</sup> | 91.3±2.2 (65.9 – 107.4) |
| Mean tree height (m)<sup>b</sup> | 29.1±1.7 (12.4 – 39.1) |
| Maximum tree height (m)<sup>b</sup> | 40.0±0.7 (32.9 – 48.8) |

<sup>a</sup> Means ± SE, range of plot-means in brackets.
<sup>b</sup> Trees with dbh ≥10 cm

### 2.3. Stand surveys and sonic tomography

Stand surveys that included the measurement of dbh and tree height (using a Vertex IV ultrasonic clinometer and T3 transponder manufactured by Haglöf, Långsele, Sweden) were conducted for all trees with a dbh ≥10 cm. To be consistent with Lutz et al. (2018), we also included all trees of the regeneration with a dbh <10 cm, but ≥1 cm, though we assumed them to be only of subordinate significance for the stand aboveground biomass estimates. In these trees, height was recorded with a measuring tape if tree height was below ca. 2 m. This way 220 trees with a dbh ≥10 cm and 457 young trees with a dbh <10 cm were surveyed. Standing deadwood with a dbh ≥10 cm and downed deadwood where a dbh ≥10 cm was assumed based on the present dimensions was recorded separately. For downed deadwood trunks that partly protruded from the sample plot, only the length of the part inside the plot was included in the plot balance.

For 112 trees, which is half of the 220 trees with a dbh ≥10 cm, internal stem decay was analyzed using sonic tomography. The 4–5 trees with the largest diameter per plot were selected for these analyses. These trees had a mean dbh (±SD) of 76±14 cm (range: 44–107 cm) and a mean height (±SD) of 35.6±4.5 m (range: 24.3–48.8 m). Sonic tomograms were recorded with a Picus 3 sonic tomograph (IML, Rostock, Germany) equipped with 12 sensors, which were evenly distributed along the stem circumference at ca. 60 cm and 160 cm height. The measuring principle of sonic tomography is based on the fact that sonic waves cross solid wood faster than decayed wood and have to bypass cavities (Arciniegas et al. 2014; Gilbert et al. 2016; Qin et al. 2018). Percentages of solid wood area in the stem cross-section were calculated with Picus Q74.2 software using the tomogram version SoT2, which produces less artefacts (and thus less cavity overestimation) than the alternative option SoT1 for marginal areas in the stem cross-section (Göcke 2017). The stem geometry of trees with an irregular shape of the bole cross-section (Fig. 1) was recorded with a Picus Caliper, because correct interpretation of the transmission time through the wood that is measured depends on the precise recording of the sensor positions (Burcham et al. 2023). The Picus software automatically corrects area proportions of differently solid vs. differently damaged or absent wood in the stem cross-section according to the geometry. All tomograms were visually checked for so-called cog wheel artefacts (Göcke 2017) that erroneously indicate a regular pattern of punctiform areas of wood decay along the stem circumference (Fig. 1h). These artefacts result from slow tangential sonic wave transmission between the tomography sensors and usually occur in stem cross-sections without other signs of damage. Damaged wood percentages that could be attributed to cog wheel artefacts were ignored in the analysis.

**Fig. 1.**
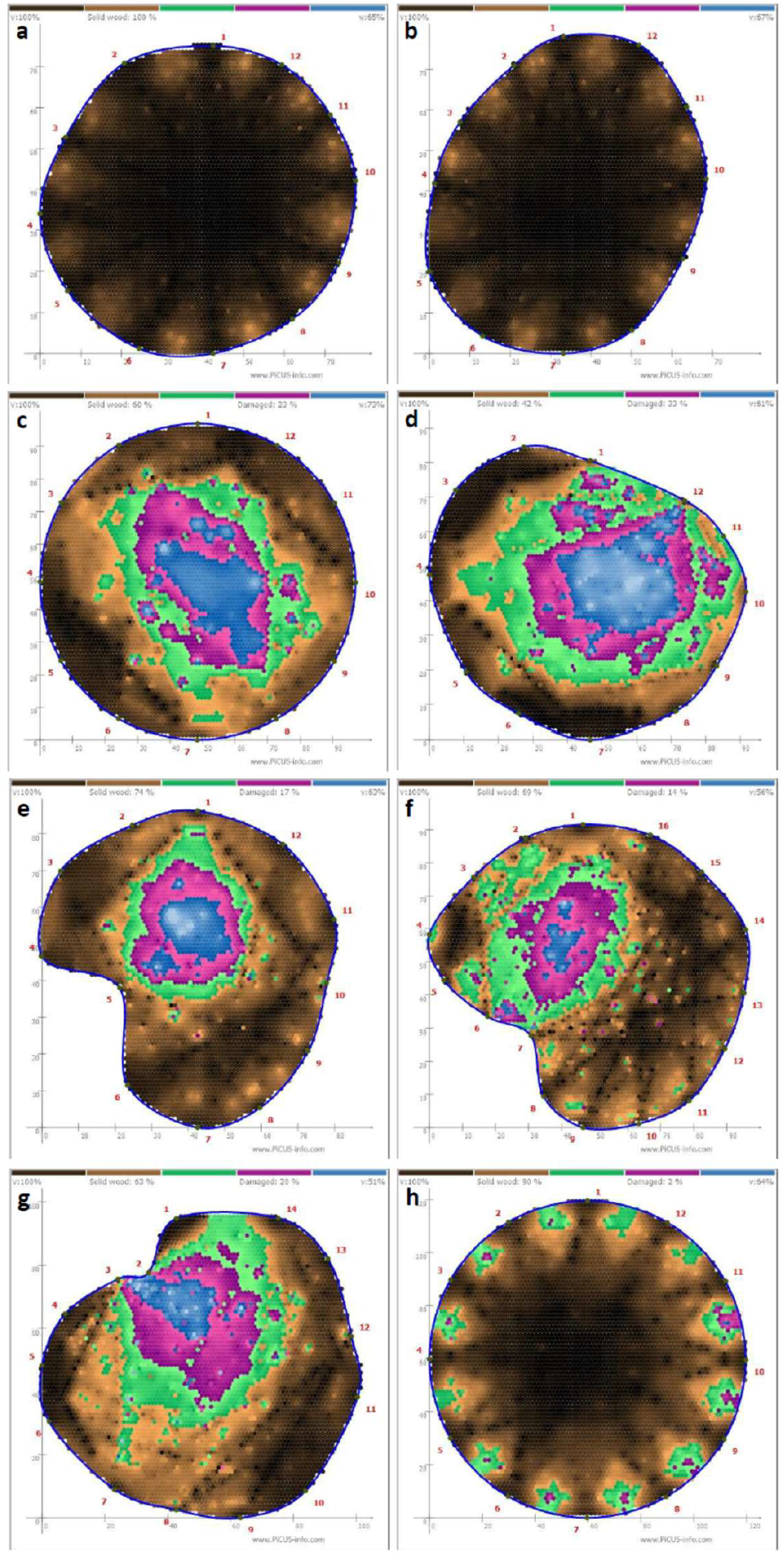
Examples of sonic tomograms of living *Pinus ponderosa* stems where brown color represents solid wood, purple color represents rotten wood, green color represents wood areas with uncertain condition, and blue color represents cavities: (a, b) stems of different shapes with completely intact wood; (c, g) stems with different extents and positions (central to lateral) of internal decay; (h) “cog wheel” artefact falsely pretending isolated lateral areas of rotten wood between sensors due to slow tangential sonic wave transmission; these artefact were corrected before calculations.

### 2.4. Allometric biomass and carbon stock estimates

Total aboveground biomass, stem biomass, branch biomass, and needle biomass were calculated for all trees of a dbh ≥1 cm using published allometric regression functions (Table S1). For *P. ponderosa*, these functions were taken from Gholz (1982), Ter-Mikaelian & Korzukhin (1997), Law et al. (2001), Jenkins et al. (2003), Tinker et al. (2010), and Vorster et al. (2020). Functions with strongly deviating results were excluded from further analyses. Stem biomass was calculated from the sum of stem wood and bark biomass, if separate equations were published. Total aboveground biomass was calculated from own published functions for aboveground biomass and by summing up mean stem, branch, and needles biomasses. The values of aboveground biomass calculated in these two alternative procedures were subsequently averaged to the final aboveground biomass estimate. The procedure for selecting allometric biomass functions was done separately for trees with a dbh >5 cm and trees with a dbh of 1–5 cm, because some biomass function, which worked well for trees of larger diameters did not work for very small tree individuals, depending on the coverage of tree sizes during the establishment of the allometric functions.

Only very trees on our sample plots did not belong to *P. ponderosa*; this concerned 4 trees of *P. contorta* and 1 individual of *Abies grandis*. For aboveground biomasses of these tree species, we used formulas of Jenkins et al. (2003). For stem, bark, and branch biomasses of *P. contorta*, we applied formulas of Vorster et al. (2020). The stem, branch, and needle biomasses of *A. grandis* were calculated as, respectively, 70%, 18%, and 11% of the total aboveground biomass. This estimate was based on available data from *A. alba* (*N*=108 trees; Hauck et al. 2025) and *A. sibirica* (*N*=41 trees; unpublished data) because of the lack of specific formulas for *A. grandis*. This seemed reasonable to us, as it only concerned 1 tree.

Deadwood biomass of *P. ponderosa* was corrected for the decline of wood density during wood decay by assuming a wood density of 250 kg m-3 instead of 380 kg m-3 (Chojnacky et al. 2014) based on similar assumptions for the decrease of wood density during decay in other conifers (Weggler et al. 2012; Köster et al. 2015; Wagner et al. 2015; Dulamsuren et al. 2016; Stakenas et al. 2020). By contrast, the organic carbon concentration in the deadwood was assumed not change significantly during decay, also based on published literature (Weggler et al. 2012; Köster et al. 2015; Wagner et al. 2015; Stakenas et al. 2020; Doraisami et al. 2026).

Conversion of biomass values to organ carbon stock densities followed values of wood carbon concentrations published by Lamlon & Savidge (2003). These values amounted to 52.47% for *P. ponderosa* and 50.32% for *P. contorta*. In the lack of a published value, the biomass of the only tree of *Abies grandis* in the dataset was converted with the value used for *P. ponderosa*.

### 2.5. Correction of biomass estimates for internal stem decay

Since fire damage and the invasion of the trees by fungi often takes place at the stem base (Gilbert et al. 2016), we assumed that internal wood decay was limited to the lower 1 m of the stem if damaged wood and tree cavities were only found at the lower measuring height of 60 cm. Otherwise, we supposed that internal wood decay has spread further up the trunk. In cases where biomass loss due to wood decay was observed at the higher measuring level, we assumed this damage for a trunk segment from 1 m height up to half the tree height. As internal stem decay especially from ground fire can be assumed to decrease with increasing tree height, we calculated with solid wood across the entire stem cross-section above this height. The sonic diagrams generated by the Picus software provide areas of intact and damaged wood and of cavities that are void of wood as area percentages in the stem cross-section. From these values, we calculated losses of wood volume in the stem segments that were based on the percent values of solid wood, before these volumetric data were converted to biomass-related values, using wood density. Values of wood density for our study tree species were taken from the literature. Wood density of *P. ponderosa* as well as the occasionally associated *P. contorta* was assumed as 380 kg m-3, whereas that of *Abies grandis* was estimated as 350 kg m-3 (Jenkins et al. 2003; Chojnacky et al. 2014).

### 2.6. Percentage of large old trees representing 50% of the stand biomass

To quantify the minimum proportion of trees that represented 50% of total aboveground stand biomass and carbon stock density (Lutz et al. 2018), we sorted the trees of the individual sample plots in a descending sequence of dbh values and identified the number of the largest trees of a plot that was needed to reach of 50% the stand aboveground biomass. To make the 25 sample plots with different tree numbers and biomass stocks comparable, the number of trees was expressed as percentages related to the total number of trees of the individual plot. We conducted this analysis for two different samples of trees: (1) all trees of a dbh ≥10 cm, because these trees were expected to contribute significantly to stand biomass and (2) and extended sample of all trees of a dbh ≥1 cm to be consistent with Lutz et al. (2018).

### 2.7. Statistical analysis

If not mentioned otherwise, arithmetic means ± standard errors (SE) are presented throughout the paper. All statistical analyses were run in R 4.4.1 software. The effect of large tree characters (maximum dbh, maximum tree height) and regeneration density on aboveground biomass was tested with generalized linear models (GLM) after scaling the predictors by subtracting the sample mean from the individual values and dividing the result by the sample’s standard deviation using the ‘scale’ function in R. The R package ‘performance’ 0.12.4 was used to check for multicollinearity during predictor selection; variance inflation factors in the used models ranged from 1.0 to 1.2. Pairwise comparison between biomass and carbon stock data were calculated with Wilcoxon rank sum tests and paired *t*-tests after checking for normal distribution with the Shapiro-Wilk test.

## 3. Results

### 3.1. Stand structure

Stand density in the studied forests was low with 219 trees ha^−1^ of a dbh ≥10 cm (Table 1). More than twice as much tree individuals were found in the regeneration with a dbh ≥1 cm, but the amount of trees varied greatly between the sample plots from 0 to 2650 tree ha^−1^ (Table 1). The low stand density of mature trees corresponded to the dominance of tall trees (up to 49 m) of large stem diameter with mean dbh of 61 cm and a mean maximum dbh per sample plot of 91 cm (Table 1). These large-diameter trees caused a high stand basal area of 57 m ha^−1^ for all trees of a dbh ≥10 cm and 58 m ha^−1^ if all trees of a dbh ≥1 cm were included (Table 1).

### 3.2. Aboveground biomass based on allometric biomass functions

The total stock density of aboveground biomass of trees with a dbh ≥10 cm amounted to 421±22 Mg ha^−1^ (Table 2). Tree individuals of the different stages of the regeneration with a dbh ≥1 cm, but <10 cm contributed only with 0.7% (or 3±1 Mg ha^−1^) to the total aboveground biomass stock density of trees with a dbh ≥1 cm of 424±22 Mg ha^−1^). The aboveground biomass was allocated with 75% to the stem, 20% to branches, and 5% to the foliage (Table 2). Standing and downed deadwood that was not included in the aboveground biomass estimate had a stock density of 7±3 Mg ha^−1^. These values were equivalent to organic carbon stock densities of 221 Mg C ha^−1^ for trees with a dbh ≥10 cm or of 222 Mg C ha^−1^ for trees with a dbh ≥1 cm in the aboveground biomass and 4 Mg C ha^−1^ in deadwood (Table 2).

**Table 2.** Aboveground biomass and carbon stock density of trees with dbh ≥10 cm and ≥1, <10 cm in *Pinus ponderosa* forests (*N*=25) based on allometric regression functions without or with correction for internal stem decay.

| | Stock density | | $P^a$ | Corrected to uncorrected ratio |
| --- | --- | --- | --- | --- |
|  | Allometry | Corrected |  |  |
| Biomass: |  |  |  |  |
| dbh $\geq 1$ cm | | | | |
| Total aboveground (Mg ha<sup>-1</sup>) | 423.6±21.9 | 417.8±20.9 | 0.04 | 99.3±0.4 |
| Stem (Mg ha<sup>-1</sup>) | 318.9±16.6 | 314.4±15.8 | 0.04 | 99.3±0.4 |
| Branches (Mg ha<sup>-1</sup>) | 85.7±4.3 | . | | |
| Foliage (Mg ha<sup>-1</sup>) | 22.6±1.0 | . | | |
| dbh $\geq 10$ cm | | | | |
| Total aboveground (Mg ha<sup>-1</sup>) | 420.6±22.0 | 414.8±21.1 | 0.04 | 98.9±0.6 |
| Stem (Mg ha<sup>-1</sup>) | 317.3±16.6 | 312.9±15.9 | 0.04 | 99.2±0.4 |
| Branches (Mg ha<sup>-1</sup>) | 84.9±4.4 | . | | . |
| Foliage (Mg ha<sup>-1</sup>) | 22.2±1.0 | . | | . |
| dbh $\geq 1$ cm, <10 cm | | | | |
| Total aboveground (Mg ha<sup>-1</sup>) | 3.0±1.1 | . | | . |
| Stem (Mg ha<sup>-1</sup>) | 1.5±0.6 | . | | . |
| Branches (Mg ha<sup>-1</sup>) | 0.8±0.3 | . | | . |
| Foliage (Mg ha<sup>-1</sup>) | 0.4±0.1 | . | | . |
| Deadwood (Mg ha<sup>-1</sup>)<sup>b</sup> | 7.2±3.3 | | | |
| Organic carbon: |  |  |  |  |
| dbh $\geq 1$ cm | | | | |
| Total aboveground (Mg C ha<sup>-1</sup>) | 222.3±11.5 | 219.2±11.0 | 0.04 | 99.3±0.4 |
| Stem (Mg C ha<sup>-1</sup>) | 167.3±8.7 | 165.0±8.3 | 0.04 | 99.3±0.4 |
| Branches (Mg C ha<sup>-1</sup>) | 45.0±2.3 | . | | . |
| Foliage (Mg C ha<sup>-1</sup>) | 11.8±0.5 | . | | . |
| dbh $\geq 10$ cm | | | | |
| Total aboveground (Mg C ha<sup>-1</sup>) | 220.7±11.6 | 217.7±11.1 | 0.04 | 98.9±0.6 |
| Stem (Mg C ha<sup>-1</sup>) | 166.5±8.7 | 166.5±8.7 | 0.04 | 99.2±0.4 |
| Branches (Mg C ha<sup>-1</sup>) | 44.5±2.3 | . | | . |
| Foliage (Mg C ha<sup>-1</sup>) | 11.7±0.5 | . | | . |
| dbh $\geq 1$ cm, <10 cm | | | | |
| Total aboveground (Mg C ha<sup>-1</sup>) | 1.6±0.6 | . | | . |
| Stem (Mg C ha<sup>-1</sup>) | 0.8±0.3 | . | | . |
| Branches (Mg C ha<sup>-1</sup>) | 0.4±0.2 | . | | . |
| Foliage (Mg C ha<sup>-1</sup>) | 0.2±0.1 | . | | . |
| Deadwood (Mg C ha<sup>-1</sup>)<sup>b</sup> | 3.8±1.2 | | | |

<sup>a</sup> One-sided paired *t*-test
<sup>b</sup> Standing and downed deadwood with a minimum dbh of 10 cm. Not included in the sums for total aboveground biomass and organic carbon stocks.

### 3.3. Biomass loss by internal stem decay

On the stand level, biomass losses due to internal stem decay were minor, yet statistically significant (Table 2). After correcting the biomass for internal decay, the aboveground biomass and carbon stock densities were 415±21 Mg ha^−1^ (or 218±11 Mg C ha^−1^) compared to 421±22 Mg ha^−1^ (or 221±12 Mg C ha^−1^). This difference was shown to be significant in a paired *t*-test (Table 2). These differences were equivalent to an overestimation by 1.1%, if the biomass data were not corrected for internal stem decay (Table 2). The inclusion of all trees with a dbh ≥1 cm in the sampled lowered this overestimation to from 1.1% to 0.7%, but the differences between the corrected and uncorrected aboveground biomass and carbon stock densities were still significant (Table 2).

### 3.4. Proportion of trees accounting for 50% of the stand biomass

The percentage of the largest trees needed for 50% of the aboveground tree biomass (and carbon stock) depended on the minimum dbh that was included in the sample (Fig. 2). The 15% largest trees of a stand accounted for 50% of the aboveground biomass, if the minimum dbh was set to 10 cm. If the sample was expanded to all trees with a dbh ≥1 cm, this value was considerably lower, as only the 2% largest trees were necessary for 50% stand aboveground biomass. These percent values are based on the intersection of the nonlinear regression lines in Fig. 2 with 50% aboveground biomass along the ordinate. Despite this strong decline from 15% to 2% by the inclusion of the tree regeneration with a dbh ≥1 cm, but

**Fig. 2.**
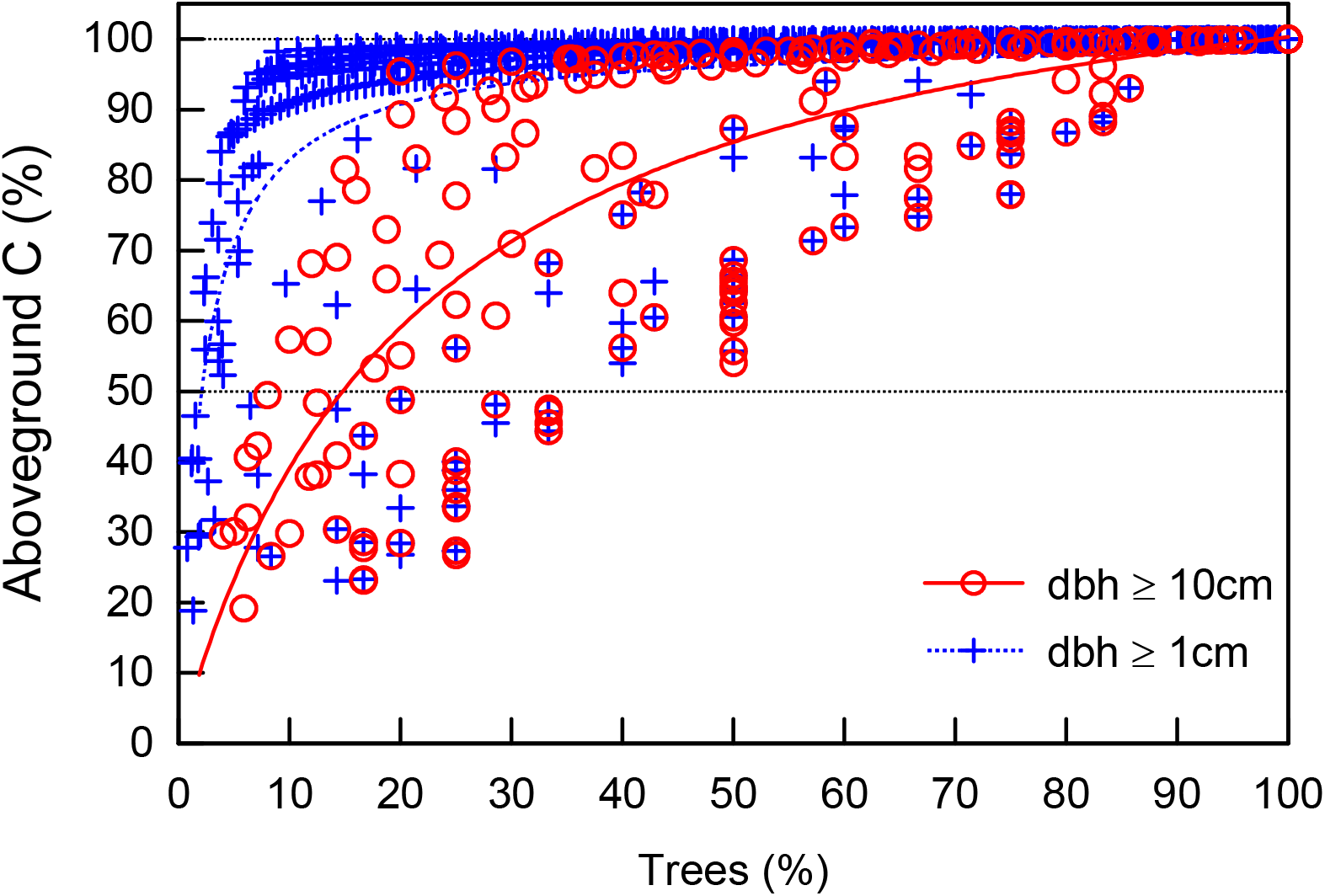
Contribution of large old trees to stand C stocks in the aboveground biomass: The diagram displays cumulative percentages of trees ordered in a sequence of descending biomass and their contributing to stand C for all trees with a dbh ≥ 10 cm (*r* = 0.78, *P*<0.001) and dbh ≥ 1 cm (*r* = 0.53, *P*<0.001). Percent values in the diagram specify the proportion of trees needed to form 50% of the stand C stock in the aboveground biomass based on the regression lines.

<10 cm in the calculation, generalized linear models showed that the regeneration densities had no significant effect on aboveground stand biomass (Table 3). This was true for both the estimates based on allometry and those corrected for biomass loss due to internal stem decay (Table 3). Rather, stand aboveground biomass was primarily determined by maximum dbh and maximum tree height and thus characters related to large trees. The strong influence of regeneration density on the percentage of trees that represent 50% of the aboveground biomass is also illustrated in Fig. 3, where this value (after ln-transformation) is taken up versus regeneration density. By increasing the total sample size, high density of trees of a dbh <10 cm strongly decreased the percent value of trees needed for 50% aboveground stand biomass (Fig. 3a). However, this parameter was not related to maximum dbh (Fig. 3b) or maximum tree height (Fig. 3c), which represented the strongest predictors of aboveground stand biomass and carbon stock density (Table 3).

**Table 3.** Generalized linear models for the effects of maximum dbh, maximum tree height and regeneration density (trees with dbh >1 cm, <10 cm) on the organic carbon stock density in the aboveground tree biomass considering all trees with dbh ≥1 cm.

| Parameter | Estimate | SE | *P* | CI |  |
| --- | --- | --- | --- | --- | --- |
|  |  |  |  | 2.5% | 97.5% |
| Allometry-based biomass: |  |  |  |  |  |
| *R*<sup>2</sup> | 0.60 |  |  |  |  |
| Intercept | 222.26 | 7.72 | <0.001 | 207.13 | 237.39 |
| Maximum dbh | 30.87 | 8.74 | 0.002 | 13.75 | 47.99 |
| Maximum tree height | 21.42 | 8.78 | 0.02 | 4.21 | 38.62 |
| Regeneration density | −0.41 | 7.97 | 0.96 | −16.02 | 15.20 |
| Biomass corrected for internal stem decay: |  |  |  |  |  |
| *R*<sup>2</sup> | 0.54 |  |  |  |  |
| Intercept | 219.24 | 7.98 | <0.001 | 203.60 | 234.87 |
| Maximum dbh | 29.71 | 9.03 | 0.004 | 12.01 | 47.41 |
| Maximum tree height | 17.19 | 9.07 | 0.07 | −0.59 | 34.97 |
| Regeneration density | 0.02 | 8.23 | 1.00 | −16.12 | 16.15 |

Model estimates (±SE) and 95% confidence intervals (CI).

**Fig. 3.**
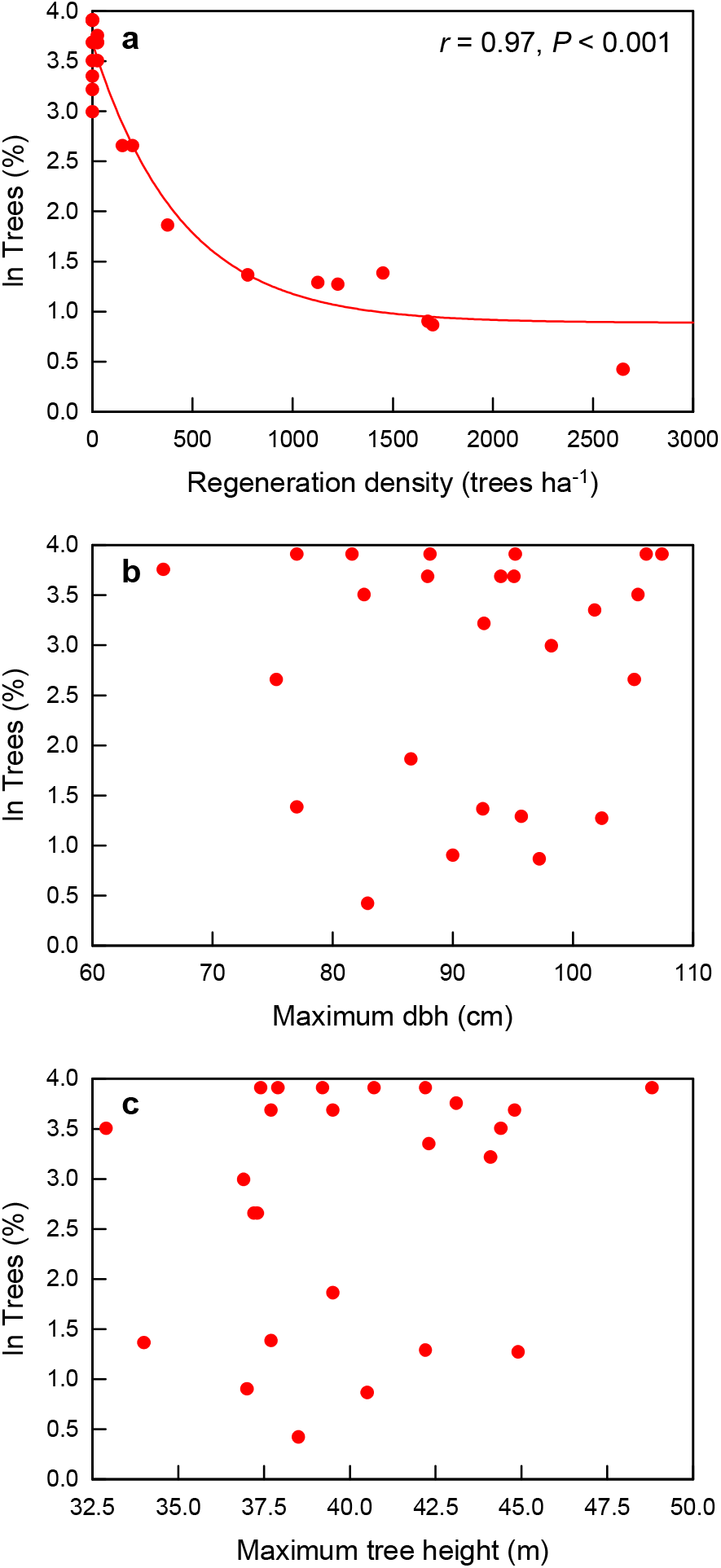
Dependence of the (logarithmized) percentage of largest trees necessary to represent 50% of the stand aboveground biomass on (a) regeneration density, (b) maximum dbh, and (c) maximum tree height of the individual forest stands (*N*=25).

In real forest stands, the biomass is, of course, not distributed among the trees in such a way that a certain number of trees accounts for exactly 50% aboveground biomass. If the number of trees was selected in our stands where the 50% biomass threshold was crossed, the realized percentages of trees needed to reach this threshold were considerably higher (Fig. 4) compared to the values deduced from the regression lines in Fig. 3. If all trees with a dbh of ≥10 cm were included, 33±3% of the largest trees were necessary to cross the 50% biomass threshold (Fig. 4a). For all trees with a dbh of ≥1 cm, this value was 26±4% (Fig. 4a). The mean percentages of stand aboveground biomass at these thresholds amounted to 58±1% (dbh ≥10 cm) and 57±1 % (dbh ≥1 cm), respectively (Fig. 4b).

**Fig. 4.**
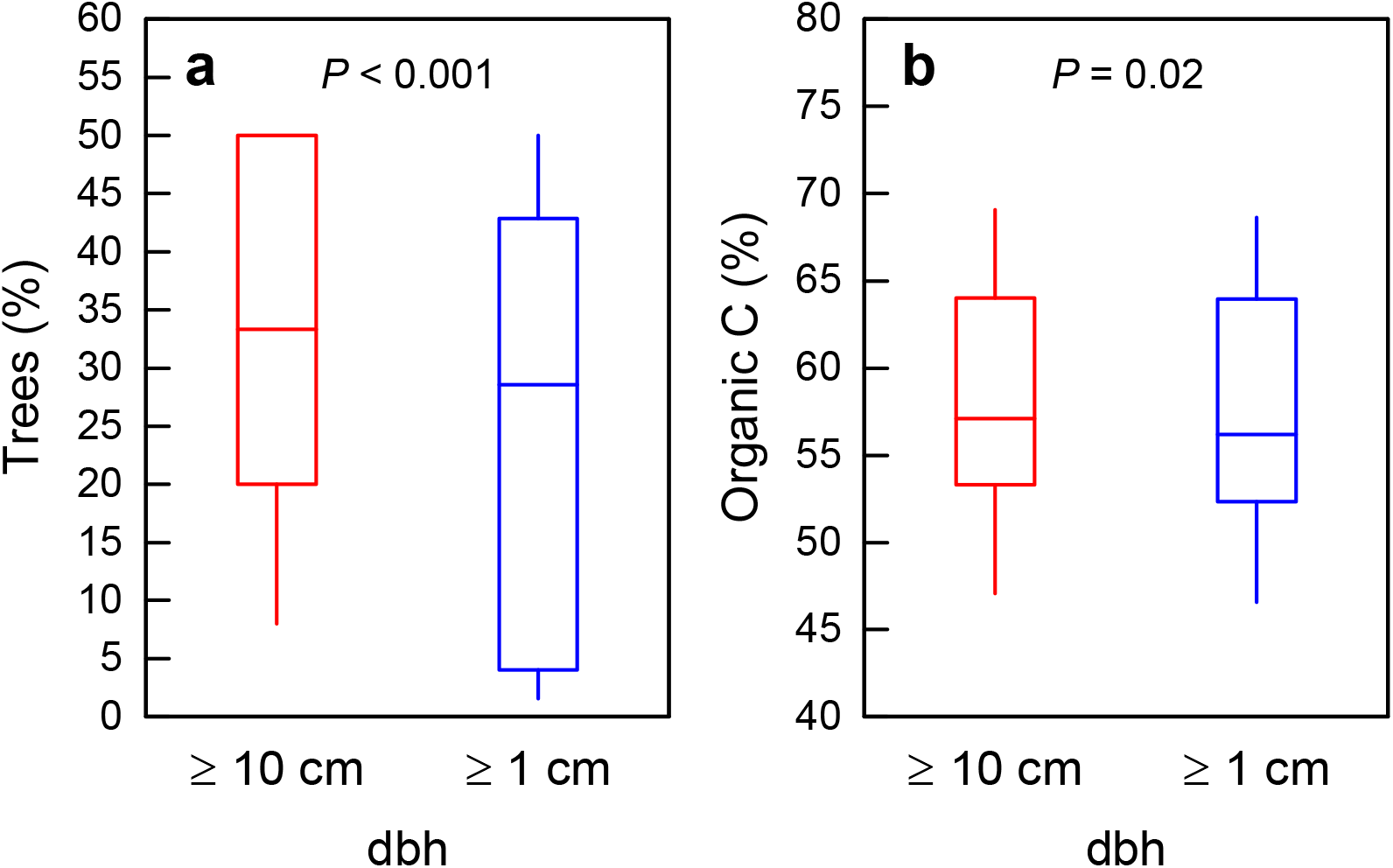
Boxplots showing the proportion of trees accounting for ca. 50% of the organic carbon in the aboveground stand biomass in *Pinus ponderosa* forests (*N*=25) considering trees with stem diameter ≥10 cm or ≥1 cm: (a) percentage of trees representing 50% of the aboveground biomass C stock, (b) the actual values of the C stock in the trees crossing the 50% stand biomass threshold. Biomass data are corrected for internal stem decay. Statistics: Wilcoxon rank sum test (two-sided).

## 4. Discussion

### 4.1 Significance of internal stem decay for biomass estimates

In the fire-prone *P. ponderosa* forests, internal stem decay and tree cavities only played a minor (albeit statistically significant) role for total aboveground biomass. The overestimation of aboveground tree biomass by 1.1% (for trees with a dbh ≥10 cm) by not correcting the allometry-based estimates for internal stem decay exceeded the corresponding value of 0.2% that was found for *Pinus sylvestris* forests in Central Europe with the same methodology (Hauck et al. 2025). Although *P. sylvestris* is a widely distributed boreal to temperate Palearctic pine species that is promoted by fire (Fernandes et al. 2008), the trees studied in the moist parts of Central Europe by Hauck et al. (2025) were definitely never exposed to wildfire. Similarly low values for biomass overestimation by neglecting internal stem decay in other Central European conifer stands not exposed to fire of 0.3% in *Abies alba* and *Picea abies* (Hauck et al. 2025) could be interpreted as an indication that the slightly higher values in the *P. ponderosa* stands of Oregon could be due to fire, especially as much of the damage visible in the field was related to the occurrence of fire scars. Yet, of course, the results from these case examples are no irrevocable proof for such a causal relationship, as our evidence is too anecdotal to ascribe such a small difference to a certain predictor. Doing this would require sampling of much more combinations of tree species and contrasting fire regimes, which would allow a statistically modeling approach.

However, what our case study can do is to substantiate that biomass loss due to internal stem decay and tree cavities in a typical temperate forest ecosystem that is frequently exposed to fire (Cova et al. 2025) is not dramatically higher than in temperate forests without wildfire. In this respect, the effect of fire in temperate forests seems to differ from termite-infested dry tropical savannas and forests, where internal stem decay was proven to be highly significant for tree biomass and carbon stock estimates (Nogueira et al. 2006; Flores-Moreno et al. 2024). In fact, stand aboveground biomass overestimation by 0.2–0.3% in fire-free coniferous ecosystems, 0.2–0.6% in fire-free broadleaved forests, and 1.1% in frequently fire-exposed coniferous forests on two continents suggest that for temperate forests, the influence of internal stem decay on biomass and carbon stock density estimates is generally minor, as has already been concluded for fire-free Central European temperate forests by Hauck et al. (2025). This confirms that generally biomass and carbon stock estimates from temperate forests, which are based on allometric biomass functions and are not corrected for internal stem decay and tree cavities, are (contrary to those from subtropical and tropical forests) valid and not significantly biased due to neglecting biomass losses in the stem interior.

While the European and western North American temperate forests differ in the relevance of fire, both are mostly managed. Hence, we have no information if the low significance of internal stem decay for stand biomass is also valid for unmanaged old-growth forests. This could differ not only because of the prevalence of older trees, but also due to potential higher diversity of wood-decomposing organisms (Majdanová et al. 2023; Yang et al. 2024).

However, old-growth stands that are unmanaged long-term are rare in the densely populated temperate climate zone at least of the northern hemisphere (Parviainen 2005; Strittholt et al. 2006; Glatthorn et al. 2018) and have thus low influence when stand-level values are upscaled to biome-wide estimates.

Published biomass values from western North American *P. ponderosa* forests vary considerably depending on stand structure, tree age and fire return intervals. Our estimates for stock densities of biomass (415 Mg ha^−1^) and carbon stock density (218 Mg C ha^−1^) in the living aboveground tree biomass exceed those of most other published values, which can be attributed to the targeted preference for stands with large-diameters trees in our study.

Working like us in central Oregon, Law et al. (2001, 2003) and Irvine et al. (2007) found organic carbon stock densities in the aboveground tree biomass of 108–129 Mg C ha^−1^ in *P. ponderosa* stands of 230–250 years. Though the stand densities in our study of 219±30 trees ha^−1^ (dbh ≥10 cm) were well comparable with mean stand densities of 234±13 trees ha^−1^ in a forest survey of 95 plots in a study by Hagmann et al. (2003) from central Oregon, stand basal area was much higher in our data (57±3 m ha^−1^) than in those of Hagmann et al. 2003 (19±1 m ha^−1^) evidencing the (intended) strong overrepresentation of large-diameter trees in our data. Studies from other areas of western North America usually investigated younger stands of *P. ponderosa*, which contained less tree biomass (Finkral & Evans 2008; Chatterjee et al. 2009).

### 4.2. The ‘1% largest trees-50% aboveground biomass’ hypothesis

Like in Hauck et al. (2023, 2025), the hypothesis that the 1% largest trees represent ca. 50% of the aboveground tree biomass (Lutz et al. 2018) could not be confirmed. This is remarkable, as this hypothesis was originally developed by Lutz et al. (2012) in temperate forests of the Pacific Northwest of western North America, though their dataset only included *P. ponderosa* as an accessory and not a dominant species like in our study. The different results in the two subsets of our data that included all trees of a dbh ≥10 cm (15%) or of a dbh ≥1 cm (2%) clearly indicated an eminent influence of tree regeneration on this parameter. Because tree regeneration only has a limited influence on stand biomass even if present at large densities (Irvine et al. 2007; Law et al. 2003; Dulamsuren et al. 2016), our results suggest that the ‘percent largest trees-50% aboveground biomass’ metrics is only meaningful if merely trees of a minimum dbh that significantly contribute to stand biomass are included in the sample. In our case, trees of a dbh ≥1 cm, but <10 cm represented only 0.7% of the aboveground stand tree biomass and were thus negligible.

The fire-prone forest ecosystems of western North America are characterized by a great variability in regeneration density depending on the point in time since the last burn and fire severity and stand structure (Hagman et al. 2013; Kolb et al. 2020; Wasserman et al. 2022). In our data, stand biomass strongly depended on attributes related to large old trees (i.e. maximum dbh and maximum tree height), but not on the density of low-diameter trees and saplings. By contrast, the ‘percent largest trees-50% aboveground biomass’ metrics was strongly influenced by regeneration density, but not by maximum dbh or maximum tree height. This supports our view that a lower dbh limit for what is accepted as a tree in this metric should be set to a much higher value than 1 cm to generate a meaningful result on the importance of large-diameter trees for conserving forest carbon. Indeed the ‘1% percent largest trees-50% aboveground biomass’ metrics of Lutz et al. (2012, 2018) has never been confirmed by other studies, though it has to be noted that large-diameter trees nevertheless remain important carbon stocks (Slik et al. 2013; Mildrexler et al. 2020; Hauck 2023, 2025).

### 4.3. Management implications

Our results confirm the high value of large old trees for organic carbon storage in temperate forests, even for an ecosystem that is frequently exposed to wildfire and thus under high risk for stem injuries. This adds to the high significance of large old trees for biodiversity conservation, which already led to application in retention forestry practices to maintain biodiversity despite management (Gustafsson et al. 2020; Emrich et al. 2025; Sillett et al. 2025). The results confirm as those of several other studies (Lutz et al. 2018; Mildrexler et al. 2020) that the retaining large old trees in managed forest stands also makes a major contribution to protect the global climate. However, not confirming ‘1% percent largest trees-50% aboveground biomass’ by Lutz et al. (2012, 2018) with our data also implies that more than 1% or very few percentages of the largest trees in a stand should be conserved to protect a major share of the stand’s organic carbon stock in the tree biomass.

## Conclusions

In contrast to our first hypothesis, frequent fire exposure in the dry temperate forests of central Oregon did not result in a major reduction of stand organic carbon stocks by internal stem decay, albeit there was a small influence detectable that was statistically significant.

Thus, termite-infested tropical savannas and some tropical rainforests remain, so far, the only known forest ecosystems with high losses of stand biomass due to internal stem decay in the living tree biomass (Nogueira et al. 2006; Flores-Moreno et al. 2024). In support of our second hypothesis and consistent with previous work in other temperate forests (Hauck et al. 2023, 2025), more than 1% of the largest trees was needed for 50% stand aboveground tree biomass, especially if the tree regeneration was excluded from the sample. However, this does not principally lower the high value of large old trees for conservation the organic carbon pool of temperate forests.

## Acknowledgements

The study could not have been carried out without permission and logistic support by the USDA Forest Service. We are thankful to Professor Klaus Puettmann (Oregon State University, Corvallis) for his help to establish contacts for field work. Felix Csapek is thanked for his help during field work.

## Authorship contributions

**Markus Hauck:** Conceptualization, Methodology, Data collection, Analysis, Writing, Editing. **Ganbaatar Batsaikhan:** Data collection, Analysis, Editing. **Germar Csapek:** Data collection, Editing. **Steffen Rust:** Methodology, Data collection, Editing. **Harold S.J. Zald:** Conceptualization, Methodology, Editing. **Choimaa Dulamsuren:** Conceptualization, Methodology, Data collection, Analysis, Writing, Editing.

