## Supplementary material for "Low impact of internal stem decay on forest carbon stocks in fire-prone *Pinus ponderosa* forests": Table S1

**Table S1.** Allometric regression functions applied for biomass estimates in *Pinus ponderosa*

| Organ | Reference | Formula[a] | a | b | Dbh[b] | |
|---|---|---|---|---|---|---|
| | | | | | > 5 cm | 1 – 5 cm |
| Stem wood | Gholz (1982) | $y=EXP(a + b\ln(D))$ | –4.491 | 2.759 | + | + |
| Stem bark | Gholz (1982) | $y=EXP(a + b\ln(D))$ | –4.206 | 2.231 | + | + |
| Stem | Law et al. (2001) | $y=a + b\,(D^2 H)$ | 4.141644 | 0.014509 | + | . |
| Stem wood | Acker in Law et al. (2001) | $y=a\,(D/100)^2 H$ | 119.33011 | | + | + |
| Stem bark | Acker in Law et al. (2001) | $y=a\,(D/100)^2 H$ | 23.115 | | + | + |
| Stem wood | Ter-Mikaelian & Korzukhin (1997) | $y=a\,D^b$ | 0.011 | 2.7587 | + | + |
| Stem bark | Ter-Mikaelian & Korzukhin (1997) | $y=a\,D^b$ | 0.0144 | 2.2312 | + | + |
| Stem | Tinker et al. (2010) | $y=a + b\,(D^2 H)$ | 4.552 | 0.019 | + | . |
| Stem wood | Vorster et al. (2020) | $y=EXP(a + b\ln(D))$ | –2.5513 | 2.2322 | + | + |
| Stem bark | Vorster et al. (2020) | $y=EXP(a + b\ln(D))$ | –3.5399 | 1.9588 | + | + |
| Live branches | Gholz (1982) | $y=EXP(a + b\ln(D))$ | –5.386 | 2.719 | . | + |
| Dead branches | Gholz (1982) | $y=EXP(a + b\ln(D))$ | –2.577 | 1.444 | . | + |
| Branches | Law et al. (2001) | $y=a + b\,(D^2 H)$ | 2.336823 | 0.007552 | + | + |
| Branches | Ter-Mikaelian & Korzukhin (1997) | $y=a\,D^b$ | 0.0469 | 2.1315 | + | + |
| Branches | Ter-Mikaelian & Korzukhin (1997) | $y=a\,D^b$ | 0.0096 | 2.4645 | + | + |
| Branches | Ter-Mikaelian & Korzukhin (1997) | $y=a\,D^b$ | 0.0045 | 2.7185 | + | + |
| Branches | Tinker et al. (2010) | $y=a + b\,(D^2 H)$ | 3.114 | 0.061 | + | + |
| Branches | Vorster et al. (2020) | $y=EXP(a + b\ln(D))$ | –5.2127 | 2.9843 | . | + |
| Needles | Gholz (1982) | $y=EXP(a + b\ln(D))$ | –4.261 | 2.097 | + | + |
| Needles | Ter-Mikaelian & Korzukhin (1997) | $y=a\,D^b$ | 0.1167 | 1.5774 | + | + |
| Needles | Ter-Mikaelian & Korzukhin (1997) | $y=a\,D^b$ | 0.0286 | 1.992 | + | + |
| Needles | Ter-Mikaelian & Korzukhin (1997) | $y=a\,D^b$ | 0.0119 | 2.0967 | + | + |
| Needles | Tinker et al. (2010) | $y=a + b\,(D^2 H)$ | 0.052 | 0.982 | + | + |
| Needles | Vorster et al. (2020) | $y=EXP(a + b\ln(D))$ | –5.75806 | 2.611 | + | + |
| Aboveground biomass[c] | Jenkins et al. (2003) | $y=EXP(a + b\ln(D))$ | –2.5356 | 2.4349 | + | + |

[a] $EXP = e^x$

[b] Application of the individual biomass functions for dbh classes: equation used (+) or not used (.)

[c] Otherwise sums of the individual organs were calculated for aboveground tree biomass
